# Novel features in the structure of P-glycoprotein (ABCB1) in the post-hydrolytic state as determined at 7.9Å resolution

**DOI:** 10.1101/308114

**Authors:** Nopnithi Thonghin, Richard F. Collins, Alessandro Barbieri, Talha Shafi, Alistair Siebert, Robert C. Ford.

## Abstract

P-glycoprotein (ABCB1) is a ATP-binding cassette transporter that plays an important role in the removal of drugs and xenobiotic compounds from the cell. It is also associated with multi-drug resistance in cancer. Here we report novel features of the cryo-EM-derived structure of P-glycoprotein in the post-hydrolytic state: The cytosolic nucleotide-binding domains (NBDs) are separated despite ADP remaining bound to the NBDs. Gaps in the TMDs that connect to the inner hydrophilic cavity are back-filled by detergent head-groups from the annular detergent micelle and are close to two regions predicted to delineate two pseudo-symmetry-related drug-binding sites. In this conformation, the (newly-resolved) N-terminal extension, NBD-TMD linker region and gap-filling detergents all appear to impede NBD dimerisation. We propose a model for the mechanism of action of the exporter where ATP will be bound to the protein for most of the time, consistent with the high physiological ATP concentrations *in vivo*.

## Introduction

P-glycoprotein (P-gp/ABCB1) is a member of the ATP-binding cassette (ABC) family of proteins(1–3). It operates as an ATP-dependent exporter in the plasma membrane and it exports a host of small, mostly hydrophobic, molecules from the cell. Its principal role is to protect against xenobiotic compounds entering the organism from the diet or environment. However in some cancers being treated by chemotherapy, the high mutation rate, combined with the strong selective pressure of the drug, frequently leads to up-regulation of P-gp and multi-drug resistance (4,5). There is therefore an unmet need for an effective and reversible P-gp inhibitor to address its mediation of multi-drug resistance in cancer. To this end, structure-informed drug design/optimisation may be useful.

There are now many atomic models and experimental density maps for P-gp deposited in the Protein Databank (PDB) and Electron Micoscopy Databank (EMDB) (6–14), and data are available at a resolution allowing a direct modeling of the amino acid residue side-chains(6–13). Murine P-gp has so far been predominant in these studies. The murine version of the protein favours an inward-facing conformation (i.e. where the transmembrane domains -TMDs- surround a cavity leading to the cytoplasm and with the nucleotide-binding domains -NBDs- separated). This configuration of the protein has been proposed to represent the higher affinity state for transported substrates (allocrites) such as drugs or xenobiotic compounds. So far, only one structure for P-gp is in the outward-facing state: The TMDs here surround a cavity leading to the extracellular milieu and the NBDs are sandwiched together, concertedly binding two ATP molecules. This structure is for human P-gp where mutation of the glutamate residues involved in ATP hydrolysis was carried out (6). The mutagenesis (E->Q) also removed two charges in negatively-charged patches which must approach each other in the ATP-bound NBD-NBD interface. The change in electrostatics may favour the formation of an NBD1-NBD2 dimer. Hence the structural and biochemical data so far imply that wild-type purified Pgp exists only transiently in the outward-facing state, even in the presence of high ATP concentrations (14,15).

Biophysical measures of wild-type P-gp dynamics have similarly demonstrated that the inward-facing configuration was dominant, also in the presence of a non-hydrolysable ATP analogue (15,16). This suggests that prevention of ATP hydrolysis alone does not favour the outward-facing state, and was consistent with the idea that electrostatic repulsion between the NBDs was more important. However these studies also found that, when trapped in a post-hydrolytic state, P-gp existed in a roughly 50:50 mixture of inward- and outward-facing conformations (16). We therefore decided to study wild-type murine P-gp under these conditions using cryo-electron microscopy and single particle analysis. Our hypothesis was that this would allow the study of both inward- and outward-facing conformations within the same population of molecules.

## Results & Discussion

### Cryo-EM

The two stages in the purification of P-gp are displayed in Supplementary Figure 1, with the final material judged to be highly pure (panel a) and monodisperse by size-exclusion chromatography (panel b). Motion-corrected electron micrographs of the vanadate-trapped P-gp also showed a homogeneous particle distribution across the electron microscopy grids, with few small aggregates (Figure 1). Particles displayed a variety of projections; side-on projections could be readily discerned, showing a distinctive annular detergent micelle surrounding the TMDs at one end of the protein (Fig. 1a, circled particles). Visual inspection of the raw particle data did not allow an unequivocal identification of P-gp in the outward-facing state, but particles in the inward-facing conformation, with the two NBDs clearly separated, were discernible in the particle population (Figure. 1a, encircled top right). After selecting particles, classification revealed well-differentiated 2D projection classes with internal features consistent with the resolution of helical secondary structural elements (Figure. 1b). Class averages displaying the inward-facing conformation were apparent whilst no outward-facing class averages were evident. Subsequent image processing of the entire particle dataset yielded three 3D classes that were very similar and all displayed an inward-facing conformation (Figure. 1c). The particle dataset corresponding to the 3D class with the highest resolution was split into two and each half was independently refined to allow for assessment of the resolution limitations of the structural analysis using the Fourier shell correlation (Supplementary Figures 2&3). A Fourier shell correlation value of 0.143 was reached at 7.9Å. The structural data obtained was therefore at *lower* resolution than was possible by cross-linking and extensive mutagenesis of the protein (6,13); however it was at *higher* resolution than data previously obtained by cryo-EM for active P-gp in the presence of a stabilising antibody fragment (14). Flexibility may be a prerequisite for active protein, hence may be an intrinsic limitation to structural studies of single particles where there are no constraints due to a crystal lattice.

**Figure 1.**
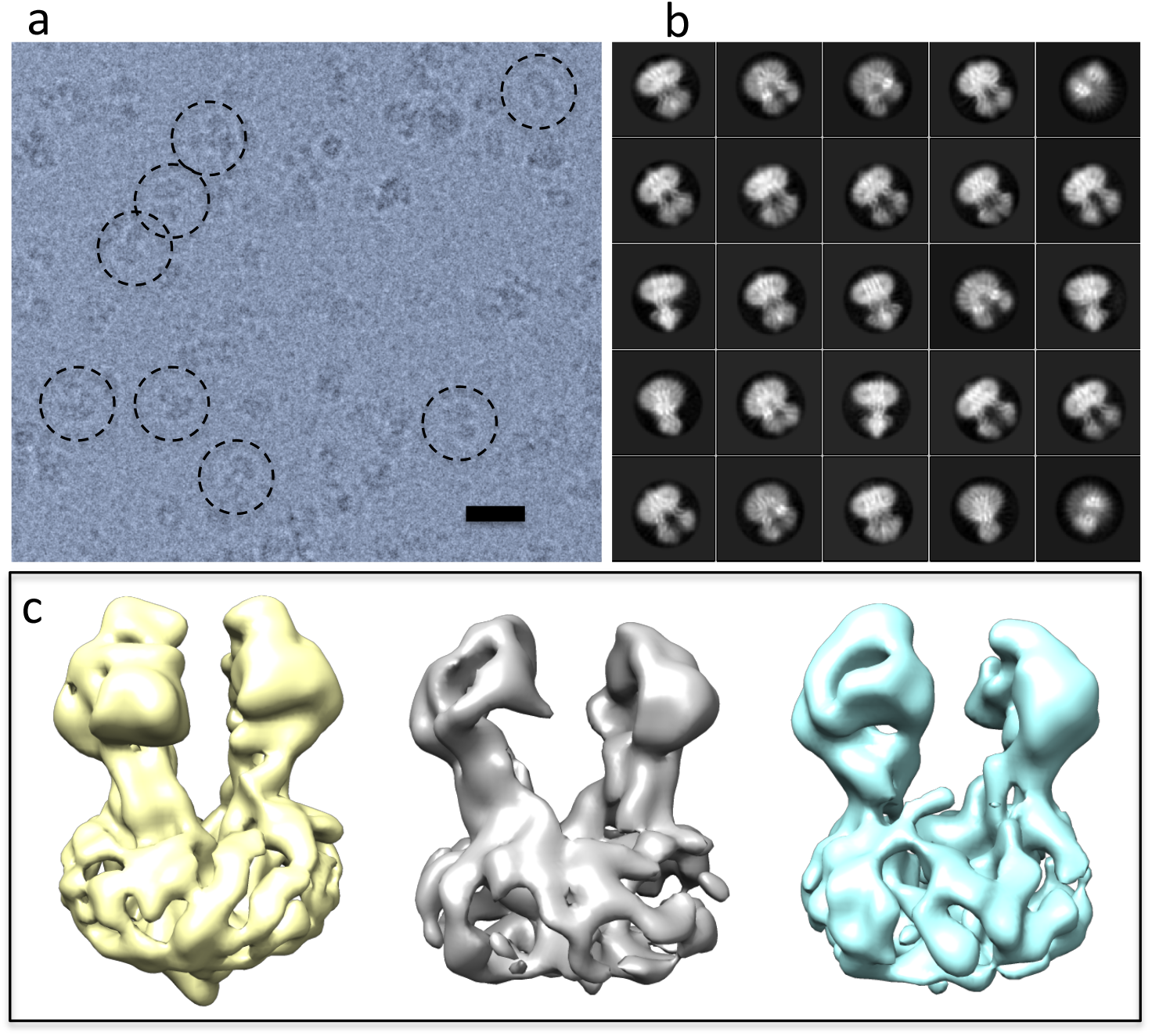
(a) Structural data. Example of a field of P-glycoprotein particles (darker grey) embedded in vitreous ice (lighter grey). Scale bar=20nm. The micrograph was filtered to 8Å resolution for clarity. Various orientations of the particles in the ice gave rise to different projections. Encircled are examples of projections corresponding to side-on views of the molecule (i.e. perpendicular to the long axis), with a clear inward-facing conformation top right, (b) Reference-free classification of the particle data set identified different projection classes and (c) 3D classification yielded 3 classes all corresponding to an inward-facing conformation of the protein.

### Interpretation of the map

Several previously-obtained mouse P-gp atomic models were tested using molecular dynamics flexible fitting (MDFF), with the inward-facing state corresponding to the PDB code *4ksb* having the best final cross-correlation coefficient with the experimental map (12). Figure 2 shows the overall 3D Coulomb density map using colours to indicate different features of the P-gp/detergent micelle complex that were interpreted after MDFF:

**Figure 2.**
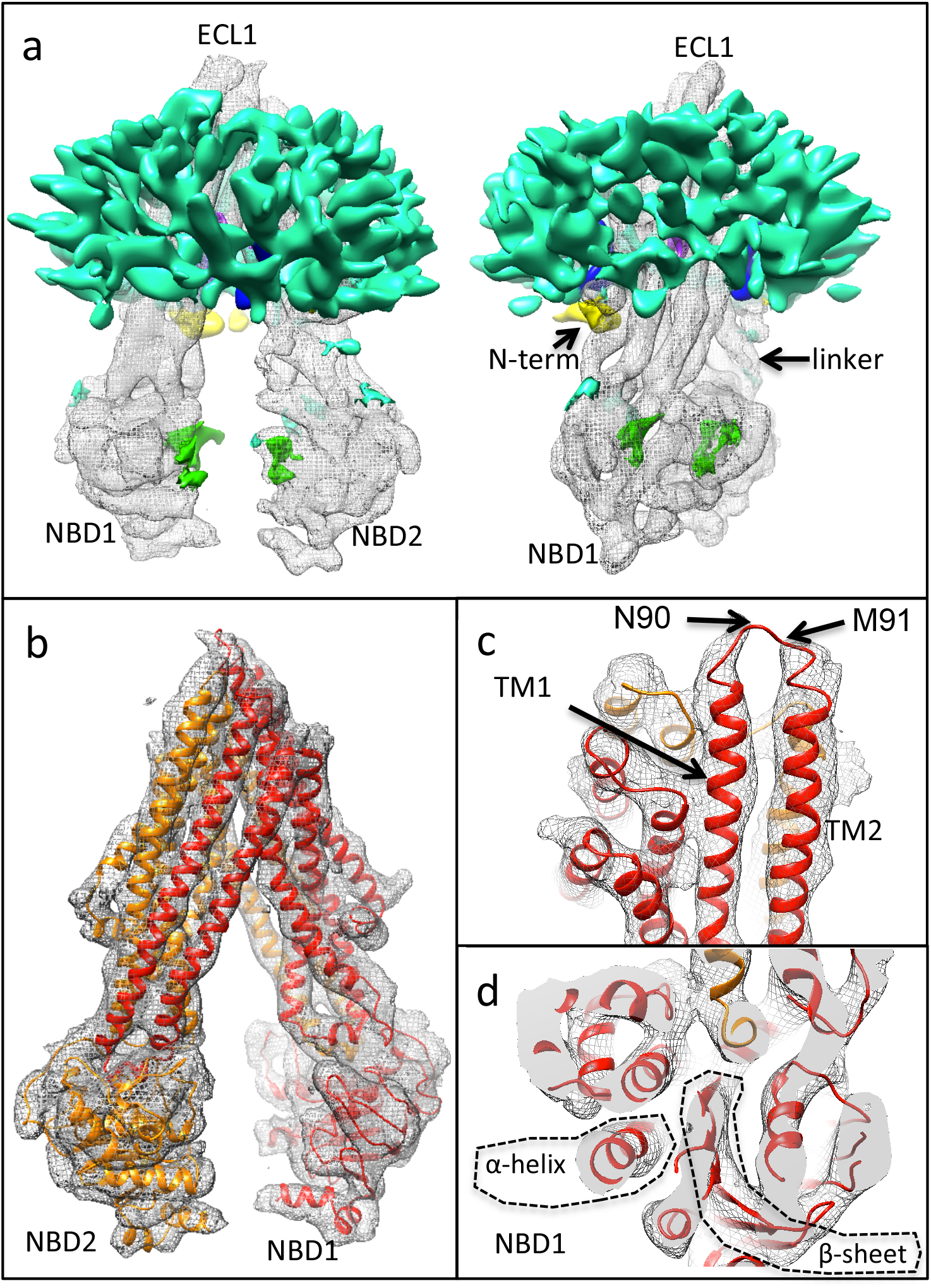
Cryo-EM map of P-glycoprotein derived from the highest resolution 3D class, (a) Orthogonal views of the map and different regions of the map are coloured according to their interpretation: Grey mesh - regions equivalent to the existing structures and within 2Å of the fitted model (4ksb after MDFF); Turquoise - detergent micelle and unassigned density; Yellow - N-terminal extension (background/left); Green - putative ADP molecules; Purple and Blue - see next Figure. The map is displayed at a threshold enclosing a volume of 75xl0^3^ Å^3^ for the core regions (grey mesh) and 85xl0^3^ Å^3^ for features not accounted for by the fitted model, (b) Core regions of the map with the MDFF-refined model shown as red and orange ribbon representations (N- and C-terminal halves of the protein, respectively), (c) Shows the extracellular portions of the map, with density lacking for 2 residues (N90, M91, arrows), (d) Shows part of NBD1. Individual β-strands viewed end-on cannot be resolved whilst individual α-helices in different orientations can be identified.

### Detergent (DDM) micelle (turquoise density)

This portion of the map in Figure. 2a is surrounding the hydrophobic membrane-spanning portions of P-gp. The maltoside polar head-group region of the detergent molecules forms this density, whilst the hydrocarbon chains of the detergent do not scatter electrons significantly more than the surrounding solvent (water). Hence the micelle appears as a hollow annular shell around the transmembrane helical portions of P-gp. A similar phenomena was reported in other cryo-EM studies of membrane proteins at resolutions exceeding 10Å. In a study examining the 3D structure of a bacterial inner membrane complex Wzz in DDM it was shown that the annular micelle could encase several independent Wzz transmembrane domains (17). The micelle surrounding P-glycoprotein does not show a smooth surface, but rather has a granular nature. This has been observed in several cryo-EM studies of membrane protein/detergent micelle complexes, and presumably results from some local organization of the detergent polar head-groups due to the high curvature of the micelle. It is likely that water molecules intercalate into this region of the micelle and account for the lower density areas in the lattice (18). Interestingly, although the P-gp transmembrane helices form a teepee-like configuration, the micelle does not conform to this, but rather displays a regular discoid shape with rough bilateral symmetry between the top and bottom halves of the annulus. However analysis of the micelle using the Resmap software (19) implies that the upper portion of the micelle is less ordered than the lower portion, which has more contact points with the protein (Supplementary Figure 3).

### P-gp core domains (grey mesh)

Figure. 2b shows the atomic model and the extracted regions of the map assigned to density within 2Å of the MDFF-refined model. Density is present for all the regions of the model except N90 & M91 in the first extracellular loop (Figure. 2c). This extracellular loop is normally glycosylated, with 3 consensus N-glycosylation sites, only one of which (N90) lies outside significant map density in the fitted model. In the *P. pastoris* expression system employed, only core glycosylation is likely to be present. In some prior structural studies, the three glycosylation sites were mutated to glutamine prior to crystallisation of the protein (7,8) implying that this loop may be flexible, variable in its glycosylation and could hinder 3D crystal formation. Although cryo-EM studies of the inward-facing conformation of ABC family members have revealed that the NBDs may be less ordered than the TMDs (20,21), in the current map, the NBDs are equally well resolved (Figure. 2d). Individual α-helices in the NBDs are observed as discrete cylindrical entities, however, individual β-strands cannot be resolved in the β-sheet displayed in Figure. 2d. These observations are consistent with the estimation of the resolution of the overall 3D map and of local regions within the map.

### Drug binding sites (Purple density)

Initial structures for P-gp with cyclic peptide inhibitors (7,8) showed an inhibitor binding site close to the apex of the internal V-shaped cavity of the TMDs. We observed a void in the equivalent region of the P-gp map (Figure 3a, dashed circle). However immediately below this, towards the cytoplasmic side of the membrane, we observed two strong densities that were not accounted for by the initially-fitted model, nor by the MDFF-refined model. Figure 3b indicates the residues predicted to be surrounding these two densities in the map. The origins of these densities remains enigmatic as there was no drug added in these structural studies. Detergent itself has been identified as a P-gp allocrite (22), hence the densities may represent the head-group regions of two DDM molecules. Higher resolution data is needed to test this hypothesis. Other studies of P-glycoprotein have aimed to identify allocrite binding sites: The Zosuquidar-bound structure of a human/mouse P-glycoprotein chimera (13) showed two drug molecules bound at the same location as the two additional densities in Figure. 3b. Compared to the map presented in Figure 3, the cavity for Zosuquidar appears to be slightly smaller - perhaps because of the cross-linking of the NBDs or the addition of the UIC2 antibody. These factors may have forced a narrower angle to be subtended between the two TMDs in the Zosuquidar study (13). Similarly, a study of P-glycoprotein with cyclic peptides designed to probe the allocrite-binding pocket showed two molecules bound in the same locations as in Figure 3. In contrast, a study of P-glycoprotein with a strong inhibitor showed its binding site to be asymmetric and much closer to the extracellular side of the molecule (10). Hence from the various structural studies so far there appear to be alternative inhibitor binding sites whilst the allocrite-binding site(s) appears to be consistently in the same location, are large, and this region in the structure has the potential to be occupied by two allocrites simultaneously.

**Figure 3.**
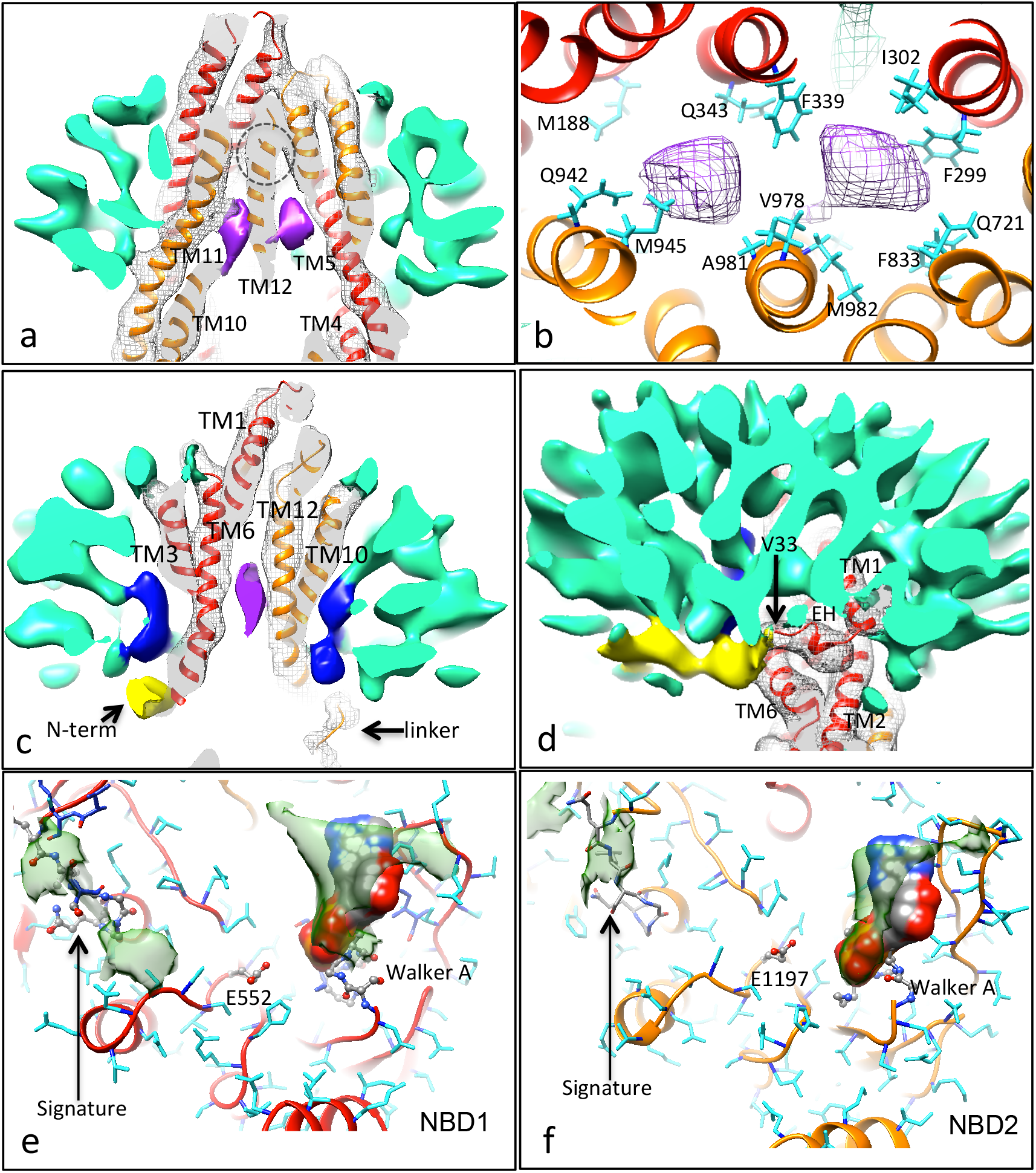
Novel features of the cryo-EM map: (a, b) Sections through the transmembrane region, (a) Shows two additional high density features (purple) that sit just below the cyclic peptide inhibitor binding site (dashed circle) and mostly formed by transmembrane (TM) helices 10,11,12,4,5 & 6 (clipped away for clarity), (b) Residues surrounding the two central densities (purple mesh) in the MDFF atomic model viewed from the extracellular side, (c) Blue density continuous with the detergent micelle (turquoise) fills the gaps formed by the inverted ‘V’ of TM helices 3&4 (left) and 9&10 (right, TM helices 4 and 9 are clipped away for clarity), (d) Additional density (yellow) below the detergent micelle (turquoise) that extends to the start of the elbow helix (EH) beginning at V33 in the fitted model (4ksb). This wraps over the protruding TM6 that connects to NBD1. (e, d) Additional density close to the interfacial surfaces of NBD1 and NBD2 (green transparent surface). In both cases elongated densities are located at the expected ADP/vanadate site. The Walker A residues and the conserved catalytic glutamates in the Walker B regions are highlighted in ball and stick representation with coloring by element. The rest of the NBD atoms are shown in stick representation with dark blue = main chain and sky blue = side chain atoms respectively and with secondary structure indicated by the ribbons. Two further small additional densities lie close to the Signature region in each case.

### TMD gaps (Blue density)

ABC transporters in the same class as P-glycoprotein display gaps between the 4^th^ and 6^th^ membrane-spanning helix in each TMD when in the inward-facing conformation. These gaps will expose the inner leaflet of the phospholipid membrane to water that fills the internal cavity created by the teepee-shape of the transporter in this conformation. Here we report L-shaped regions of density protruding into these gaps from the lower surface of the detergent micelle (Figure 3c). These densities appear to be due to distortion of the micelle in this region, with the hypothesis that maltoside head-groups kink inwards to fill the gaps that would otherwise lead to the exposure of detergent hydrocarbon chains to the aqueous central vestibule. It is possible that, i*n-vivo,* inner lipid leaflet-located molecules may behave similarly, allowing access to the transporter via these two portals. In support of this interpretation, studies have shown that detergent molecules such as DDM can operate as P-gp substrates at low concentrations (22).

### N-terminal extension and linker region (yellow density)

Structural data for the highly charged N-terminal ~30 residues of P-gp is lacking so far, however a 30Å-long extended density that joins with the N-terminal ‘elbow’ helix of P-gp can be observed in the map (Figure. 3d). This region has a finger-like shape and the fingertip merges with the head-group region of the detergent micelle. If this kinked density were to be fitted with 3 short α-helices with two breaks, it would account for most of the remaining ~30 residues of the P-gp N-terminus. Unambiguous assignment of the secondary structure in this region is stymied by the current resolution limitations, however. If this region in the protein behaves similarly *in-vivo*, then it will likely interact with the polar head-group region of the plasma membrane and would have to move from this position in order to allow NBD dimerization and the formation of the outward-facing state.

Little structural data exists for the P-gp linker region (residues 627-691) that joins the end of NBD1 with the elbow helix of TMD2 (residues 692-702). Here, in common with the 4ksb atomic model residues 684-691, we observed a 25Å-long curved density (foreground and right, Figure. 2a) that connects with the TMD2 elbow helix, and which wraps around the surface of the cytoplasmic portion of TMD2 helix 12. This additional density is initially manifested in the map close to the start of TMD helix 9 from a position that is juxtaposed between the two NBDs. This implies that the C-terminal end of the linker region is capable of being ordered under certain conditions, and (akin to CFTR/ABCC7 in the inward-facing conformation (20,21)), may play a role in regulation of NBD dimerisation.

### Nucleotide binding (Green density)

Two additional weak regions of density lie outside the NBDs but in the expected positions for nucleotides (Figure. 2a). These are probably due to ADP and vanadate. The density close to NBD1 (Fig. 3e) displays higher density than that at NBD2 (Fig 3f), implying that the NBD1 site may have a higher occupancy. Several reports suggest that NBD1 has a higher affinity for ATP and ADP/vanadate than its C-terminal counterpart (9,23,24). We note that some weak additional density is found near the Signature regions and between the NBDs and (Figure. 3e, f). It seems possible that this could represent some local organization of the linker region (see above).

The identification of significant regions in the map not accounted for by the atomic model was checked for model- and fitting-independence. This was done by comparing the results described above with those obtained by a very simple rigid body fitting of the first-half and second-half structural units of the atomic models (Supplementary Figure 4). Even with this basic fitting procedure, all the above-mentioned additional density regions were apparent.

### Implications of the P-gp structure for mechanistic models

The current structural data implies that the ADP/vanadate state (post-hydrolytic state) can be predominantly inward-facing with the NBDs separated. This is somewhat different to conclusions from a recent electron paramagnetic resonance (EPR) study of spin-labeled cysteine residues that suggested that about half the P-gp molecules would be in the outward-facing state in the presence of vanadate and nucleotide (16). A prior cryo-EM study of vanadate-trapped P-gp at low resolution (EMD-3427) also showed an inward-facing configuration (14), although with the caveat that the conformation was stabilized by a conformation-specific antibody. Nevertheless, the NBD separation and overall configuration of P-glycoprotein shows a close correlation between the current 8Å-resolution map and the 20Å-resolution map with F_AB_ bound (Supplementary Figure 5).

A simple scheme incorporating the switching between the inward and outward-facing states is shown in Figure 4: In this model, the inward-facing, ATP-bound conformation of P-gp predominates at the high ATP levels in the cell (equilibrium at stage 1). Formation of the outward-facing state may be disfavored because of electrostatic repulsion in the NBD dimer (red circle) combined with the need for removal of steric blockers (the NBD1-TMD2 linker region and the polar head-groups of lipids filling the TMD gaps). These various properties of the exporter could prevent unwanted ATP hydrolysis under normal physiological conditions where no allocrite was present. Formation of the outward-facing state in the absence of allocrite is therefore probably a rare event and will determine the basal ATP hydrolysis rate. When a P-gp allocrite is bound (green hexagon), the equilibrium at stage 1 is shifted, and formation of the outward-facing state becomes less rare. Upon formation of the outward-facing state, the escape of allocrite to the outside of the cell and ATPase activity can occur (followed by a rapid reversal to the inward-facing, post-hydrolytic state - stage 2). Inorganic phosphate and ADP must then dissociate to form the apostate (stage 3) before ATP can re-bind (equilibrium 4). There is no obvious reason why allocrite should not bind to any of the inward-facing states detected so far in structural studies. Access to the two proposed allocrite-binding sites may be via the nearby gaps between transmembrane helices 4&6 and 10&12. If trapped by vanadate, then the protein will remain in the inward-facing state with no further transport, nor ATP binding and hydrolysis. Stabilisation of the outward-facing state may only be possible by mutation of certain residues at the NBD interface (such as the Walker B motif Glutamate residues) or by crosslinking the NBDs when they are transiently together.

**Figure 4.**
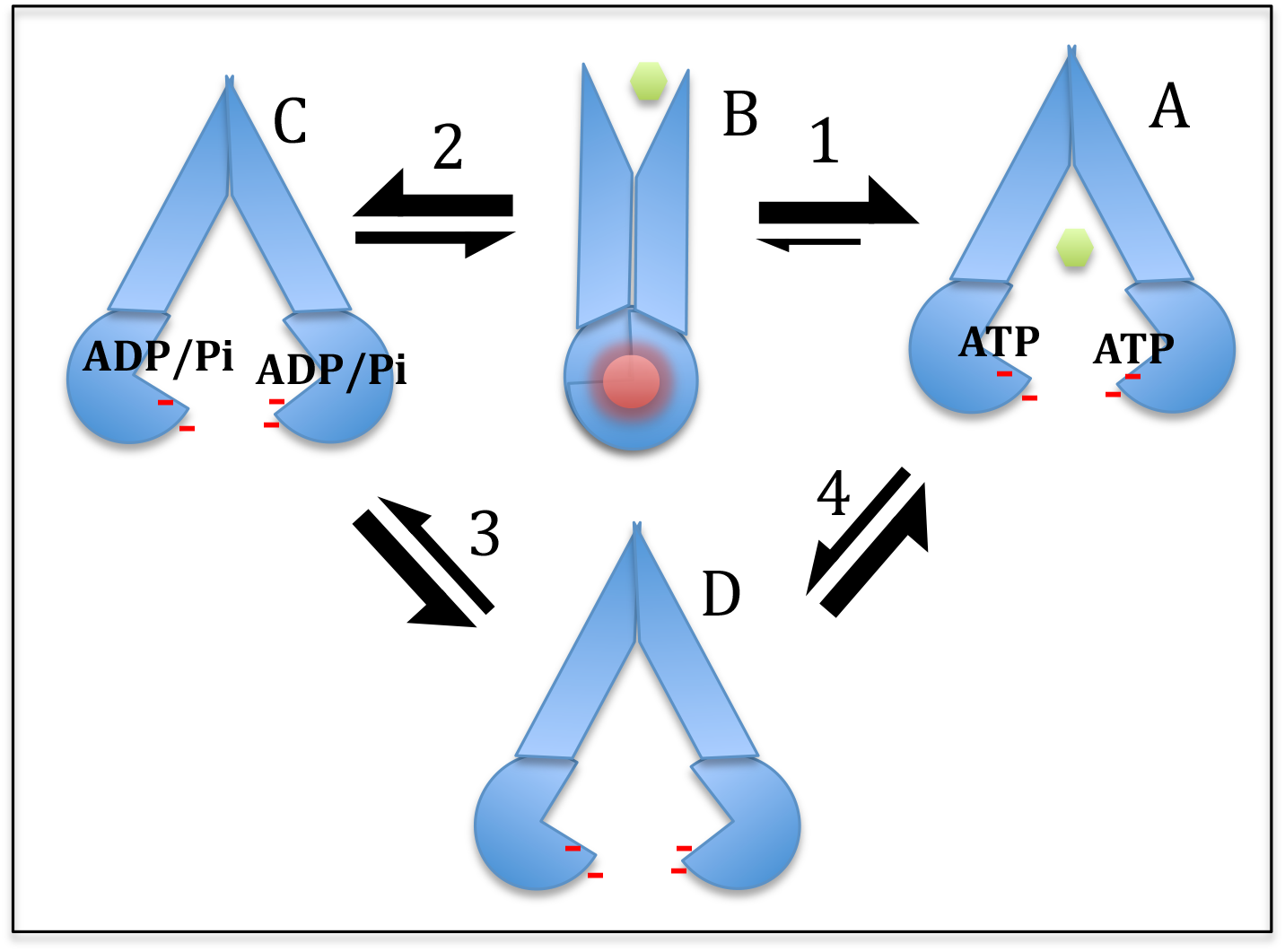
Model of P-glycoprotein action. Under physiological conditions (high ATP, no allocrite) the inward-facing state (A) predominates over the outward-facing state (B) which is unstable because of bringing together negative charges (red dashes, red circle). Allocrite binding (orange hexagon) increases the chances of formation of the outward facing state and export of the allocrite to the outside. Hydrolysis of ATP can occur during the lifetime of the outward-facing state and hence there is a finite chance of forming the post-hydrolytic state (C), even in the absence of allocrite. Dissociation of inorganic phosphate (Pi) and ADP results in the formation of the transient apo-state (inward-facing, D) before ATP re-binds and returns the system to the starting point (A). The thickness of the two-way arrows at stages 1-4 indicate the relative proportions of the different paired states, but are not intended to be quantitative.

Whilst satisfying most of the structural and biophysical data on purified P-glycoprotein, the model in Figure 4 does not concur with the more generally-accepted idea that *ATP binding* to P-glycoprotein and similar ABC exporters *drives* dimerization of the NBDs and the switch to the outward-facing state with concomintant export (25,26). One might reject the implications of the published structural and biophysical studies because they represent isolated systems. The importance of the lipid bilayer, for example, may be crucial (27–30). Secondly, it is possible that the phosphorylation state of the NBD1-TMD2 linker region may be involved in regulating the switch from inward-to outward-facing states(31), as implied for CFTR/ABCC7 (20). However if one changes the word *‘drives’* to the four words *‘increases the chance of’* in the first sentence of this paragraph, then the models for P-glycoprotein mechanism become more compatible. The model derived from the structural and biophysical data would propose the need for the combined presence of ATP *and* allocrite in P-glycoprotein to sufficiently favour the formation of a (still rare) outward-facing state. Depending on the allocrite, one observes increases in ATPase activity by P-glycoprotein of 2-16 fold (11,29,32), which implies that there is ample capacity to readjust the imbalance between inward- and outward-facing states in the system.

## Methods

### Materials

Yeast growth media was purchased from Formedium (Hunstanton, Norfolk, UK). Reagents for protein purification, sodium orthovanadate and disodium ATP were from Sigma-Aldrich (Dorset, UK). n-Dodecyl-β-Maltoside detergent (DDM) was from Merck Chemicals (Nottingham, UK). HisTrap HP and Superose 6 columns were from GE Healthcare (Buckinghamshire, UK). Quantifoil R 1.2/1.3 grid was purchased from Agar Scientific (Essex, UK).

### Mouse P-gp expression in Pichia pastoris

Cells containing the *opti-mdr3* gene used in this study were kindly provided by Prof. Ina L. Urbatsch, Texas Tech University (33). Cell culture was conducted using a previously described shake-flask method with minor modifications (34). Protein expression was initiated by an addition of 0.5% (v/v) methanol and boosted with an equivalent addition of methanol at 24 and 48 hr after the first induction.

### Protein purification

Cell rupture and microsome preparation were described previously (35). Microsomes were diluted to 2.5 mg/ml prior to solubilisation in detergent-containing buffer (50 mM Tris pH 8.0, 10% (v/v) glycerol, 50 mM NaCl, 1 mM 2-mercaptoethanol and 2% (w/v) DDM). Protein purification via immobilized metal affinity chromatography (IMAC) and size-exclusion chromatography SEC were previously described (36) with slight modifications as follows: unbound proteins in IMAC were washed twice with buffers containing 20 and 80 mM imidazole while His-tagged murine P-gp was eluted with 200 mM imidazole. Size-exclusion chromatography (SEC) was conducted using a Superose 6 column (Supplementary Figure 1). Both steps were carried out in the presence of 0.1% (w/v) DDM. Purified protein was concentrated to 10 mg/ml using a 100 kDa cut-off Vivaspin concentrator, flash frozen in liquid nitrogen and stored at −80 °C.

### Cryo-electron microscopy

P-gp was incubated with 2 mM Na_2_ATP, 2 mM MgCl_2_ and 1 mM Sodium Orthovanadate for 15 minutes at 37 °C prior to deposition onto grids as described in (14). Quantifoil 200 or 400 Au grids with a R1.3 spacing pattern where washed multiple times with chloroform in a glass dish on filter paper to remove hydrophobic contaminants residual from the manufacturing process and left to air dry for several minutes. Grids were then placed on a parafilm-wrapped glass slide before glow discharging for 2 minutes at 25 mA. Samples in vitreous ice were prepared using a FEI Vitrobot MkIV - 3μl of protein sample at a concentration of 1mg/ml was gently placed in the middle of the grid before immediately blotting for 4 seconds and flash freezing in liquid ethane. Grids were assessed for ice thickness and specimen quality using a Polara G30 before shipping to the eBIC UK National facility for high resolution data acquisition on a FEI Titan Krios G2 microscope. Data were recorded at 300 KeV using a 20 mV energy filter and a Gatan K2 electron detector - images were recorded using a total dose of ~70 e^-^ over 40 frames at a calibrated sampling increment of 1.06 Å/pixel. Data were recorded using a defocus range between −1 and −4 μm sampled at 0.25μm increments. 2209 movie stacks were recorded and drift was removed and dose weighted using MotionCorr 2 (37).

### Image processing

A general workflow of the process is explained in Supplementary Figure 2. In brief, Images were CTF corrected using gCTF (38) and over 80% of the recorded data had resolution extending between 4-7Å - the remaining images were discarded. A random selection of 30 images was used for initial autopicking reference particles using gAutomatch (developed by Zhang K, MRC Laboratory of Molecular Biology, Cambridge, UK) and a small dataset of 4000 particles was processed to provide picking templates. gAutomatch was then used to pick an initial first pass selection of ~580,000 particles. Data was subsequently processed in RELION (39)- following CTF correction as above, and multiple rounds of 2D classification were performed to remove non-P-gp particles. Examination of the 2D classes suggested that P-gp was in an inward-facing conformation. A low-resolution start model was generated from classes representing different Euler angles and then 3D classification subdivided the dataset into three 3D classes with one class demonstrating a higher resolution than the other two. This class was subjected to auto refinement, producing a final 3D map with an estimated ‘gold standard’ resolution assessment of 7.9Å (40,41). The map was post-processed using a mask designed to fit locally around the protein density with a 6 pixel soft fade and automatic b-factor detection. Local resolution in the map was assessed using the Resmap software (19).

### Model building and molecular dynamics flexible fitting

Initially, rigid-body fitting into the cryo-EM map was performed using the UCSF Chimera software ‘fit-in-map’ routine (42). The VMD software (43) was then employed to prepare the system and to generate all the necessary configuration files for the simulation. The system was subjected to flexible fitting using the Molecular Dynamics Flexible Fitting (MDFF) package (44). A scaling factor of 0.3 kcal/mol was applied to the entire model, representing the force applied to the atoms in order to flexibly fit to the cryo-EM map with a density threshold ϕ_*thr*_ of zero (default value, corresponding to the solvent peak). To maintain the integrity of the secondary structure elements and prevent over-fitting, harmonic restraints were also applied. A simulation run was performed using the NAMD 2.12 software (45) and the CHARMM36 force field (46). The system was subjected to 100,000 steps (100ps) of energy minimization, followed by 5,000,000 steps (5ns) of the production run, without symmetry restraints, in a vacuum environment and at a constant temperature of 300K. UCSF Chimera(42) was used to visualize and analyze trajectories and finally to compare cross-correlation coefficients at defined time steps, corresponding to models extracted at different frames (Supplementary Figure 6).

The MDFF identified significant additional density compared to the fitted model. It was possible that this additional density might influence the MDFF, giving a distorted model. This was checked by doing a simple rigid-body fitting of the first structural half of the molecule (TM1-3,6,10,11,NBD1) followed by the second half (TM7-9,12,4,5,NBD2) for different atomic models.

The experimental map and MDFF-refined atomic model can be downloaded via the electron microscopy database (EMDB) under the codes *EMD-4391, PDB ID 6GDI*.

## Acknowledgements

We wish to thank Dr Swathi Lingam (University of Oxford), Dr Stephen Prince (University of Manchester) and Dr Hao Fan (ASTAR Bioinformatics Institute, Singapore) for useful discussions. We thank Dr Ina Urbatsch (Texas Tech University, Lubbock) for the P-glycoprotein-expressing *Pichia pastoris* cells. NT is supported by the Development and Promotion of Science and Technology Talent Project (DPST) and the Institute for the Promotion of Teaching Science and Technology (IPST), Thailand. AB is supported by a University of Manchester/ASTAR Singapore PhD studentship. TS is supported by the Punjab education endowment fund (PEEF) with a Shahbaz Sharif Merit Scholarship (SSMS). Data collection time at the UK cryo-EM eBIC facility was via Rapid Access mode and the Manchester Block-Allocation Grouping.

